# Viscoelasticity of biomolecular condensates conforms to the Jeffreys model

**DOI:** 10.1101/2020.11.27.401653

**Authors:** Huan-Xiang Zhou

**Affiliations:** Department of Chemistry and Department of Physics, University of Illinois at Chicago, Chicago, Illinois 60607, United States

## Abstract

Biomolecular condensates, largely by virtue of their material properties, are revolutionizing biology, and yet physical understanding of these properties is lagging. Here I show that the viscoelasticity of condensates can be captured by a simple model, comprising a component where shear relaxation is an exponential function of time and a component that is purely viscous (corresponding to instantaneous shear relaxation). Modulation of intermolecular interactions, e.g., by adding salt, can disparately affect the two components, such that the exponentially-relaxing component may dominate at low salt whereas the purely viscous component may dominate at high salt. Condensates have a tendency to fuse, with the dynamics accelerated by surface tension and impeded by viscosity. For fast-fusion condensates, shear relaxation may become rate-limiting. These insights help narrow the gap in understanding between the biology and physics of biomolecular condensates.

---

Biomolecular condensates formed by phase separation are now recognized for mediating numerous cellular processes such as ribosome biogenesis^1^ and implicated in neurodegeneration and cancer.^2^ Condensates often appear as micro-sized droplets under a microscope. Much attention has been paid to the factors that determine the phase equilibrium.^3–9^ Material properties, in particular surface tension and viscoelasticity, of condensates dictate their spatial organization,^1, 8^ shape recovery,^10^ and fusion^1, 10–12^ The first set of viscoelasticity data has recently been reported for PGL-3 protein droplets.^13^ The corrected analysis of those data has been published.^14^ Here I show that the Jeffreys model of linear viscoelasticity can be used to fit the data of PGL-3 and other biomolecular droplets. This finding makes it possible to not only resolve practical problems in the extraction of viscoelastic properties from experimental data but also gain deeper physical understanding of biomolecular condensates.

Viscosity contributes to stress, i.e., force per unit area, between different layers of a fluid. For a Newtonian fluid, this stress, 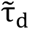, is proportional to the symmetrized strain-rate tensor 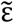:

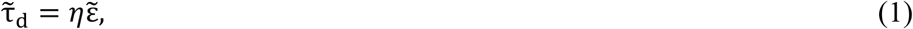

where η is the viscosity and

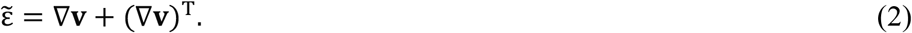

In the last equation, **v** is the velocity field in the fluid, ∇ denotes the gradient operator, and the subscript “T” denotes transpose. The hydrostatic pressure, *p*, also contributes to the stress, and hence the total stress tensor is

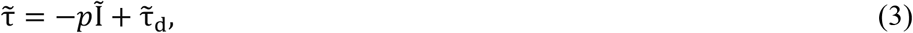

where 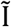 is the unit tensor. Instead of responding instantaneously to the strain rate as in a Newtonian fluid, the stress in biomolecular condensates and other complex materials depends on the entire history of the strain rate. Limiting to small strain rates so that the relation between 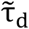 and 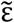 is still linear, Eq. (1) is generalized to

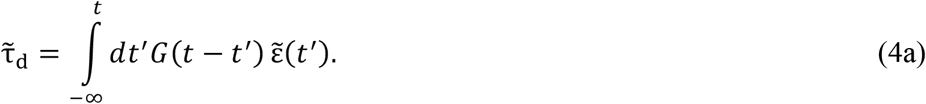

The function *G*(*t*) introduced above is the shear relaxation modulus. Note that 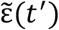 at *t*′ > *t* cannot have any effect on 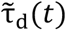. Hence we must stipulate that

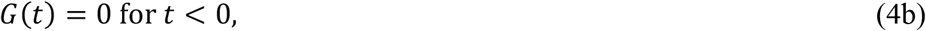

Correspondingly we can change the upper limit of the integral in Eq. (4a) to +∞,

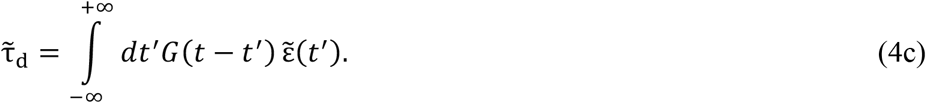

The constitutive relation, Eq. (1), for a Newtonian fluid corresponds to a delta function for the shear relaxation modulus:

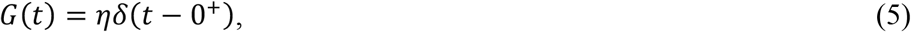

where 0^+^ indicates a peak position that is beyond *t* = 0 by an arbitrarily small positive amount.

Analogous to Eq (2) for the strain-rate tensor, we can define the shear-strain tensor

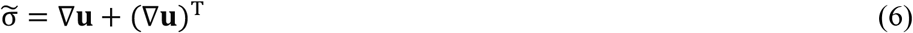

in terms of the displacement field **u**. It is obvious that

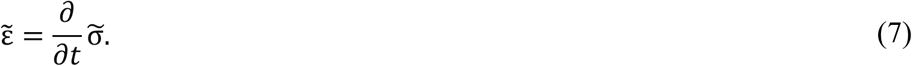

Now consider a shear strain with a sinusoidal dependence on time,

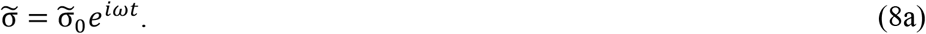

The corresponding strain-rate tensor is

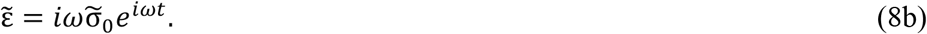

Substituting Eq. (8b) into Eq. (4c), we find

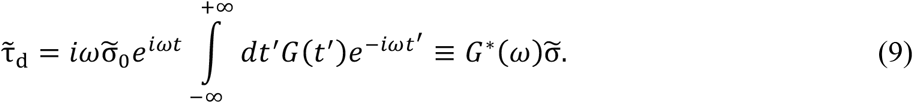

The last identity defines the complex shear modulus *G*\*(*ω*), which is related to the Fourier transform of the shear relaxation modulus,

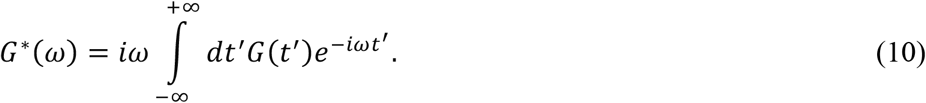

We denote the real and imaginary parts as *G*′(*ω*) and *G*″(*ω*), i.e.,

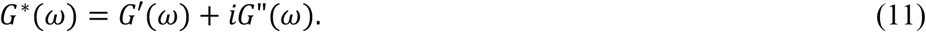

*G*′(*ω*) is called the elastic (or storage) modulus whereas *G*″(*ω*) the viscous (or loss) modulus. A Newtonian fluid [with *G*(*t*) given by Eq. (5)] has

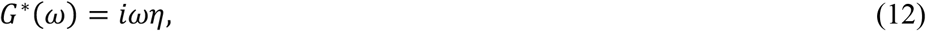

with a purely viscous modulus that is proportional to the angular frequency.

In the so-called Maxwell model, shear relaxation is an exponential function of time, with a time constant *τ*_R_,

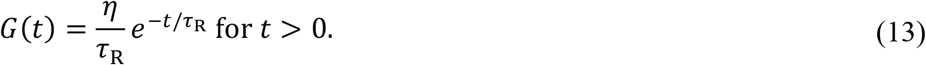

The corresponding complex shear modulus is

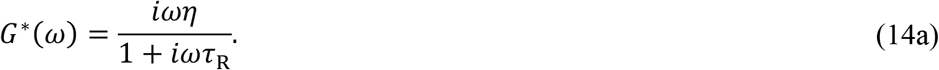

In the limit *τ*_R_ → 0, the Maxwell model reduces to a Newtonian fluid, consistent with instantaneous shear relaxation. The elastic and viscous moduli of the Maxwell model are

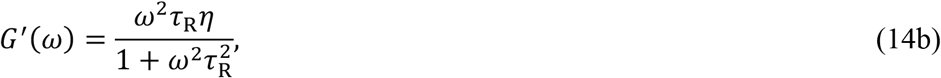

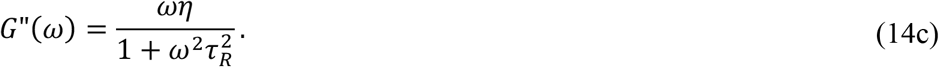

In this case, *G*′(*ω*) is a monotonically increasing function of *ω*; in contrast, *G*″(*ω*) reaches maximum at *ωτ*_R_ = 1. *G*″(*ω*) is larger than *G*′(*ω*) at *ωτ*_R_ < 1 whereas the opposite is true at *ωτ*_R_ > 1. Thus the Maxwell model behaves as a viscous liquid at low frequencies (or for slow motions) but as an elastic solid at high frequencies (or for fast motions). A linear combination of a Newtonian fluid and the Maxwell model,

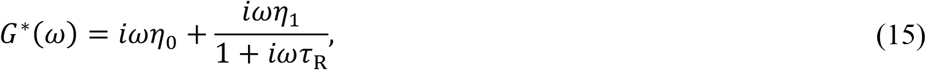

is the Jeffreys model. Dilute polymer solutions conform to this model, where the first term represents the viscous solvent, and the second term represents the contribution of the polymer solute. A main aim of this paper is to demonstrate that the Jeffreys model can be used to fit the viscoelasticity data of dense biomolecular condensates.

The first set of such viscoelasticity data was reported by Jawerth et al.,^13^ who used optical tweezers to deform PGL-3 droplets over a range of frequencies and determine the effective spring constant *χ*(*ω*) of the droplets (Fig. 1). The experiment involves optically trapping two beads (with radius *a*) at the opposite poles of an otherwise spherical droplet (with radius *R*). To analyze their data, Jawerth et al. modeled the droplet deformation dynamics by the Navier-Stokes equations, thereby expressing *χ*(*ω*) in terms of *G*\*(*ω*) and the surface tension, *γ*, of the droplet. The expression contains an infinite sum, which Jawerth et al. truncated at the 5th term because their expressions for the terms grow rapidly in complexity. I have obtained a compact expression for *χ*(*ω*) that allows for the summation to be evaluated to any order:^14^

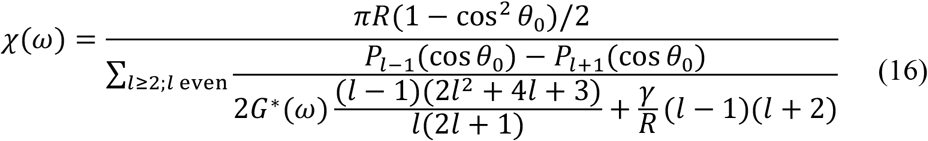

where θ_0_ = *a*/*R* and is small, at 0.1 or less, and *P*_*l*_(*x*) are Legendre polynomials.^15^ This summation turns out to converge very slowly, requiring *l* up to 40,000 to 100,000. Truncation at the 5th term (i.e., *l* = 10) thus incurs severe numerical errors, and leads to qualitatively incorrect asymptotic formulas at θ_0_ → 0. In addition to the full expression for *χ*(*ω*), Ref. 14 also presented a numerical procedure for the complete extraction of *G*\*(*ω*) from *χ*(*ω*) data, which is essentially an inversion of Eq. (16). The extraction procedure was illustrated using simulated *χ*(*ω*) data, generated by using a Jeffreys model for *G*\*(*ω*) in Eq. (16).

**FIG. 1.**
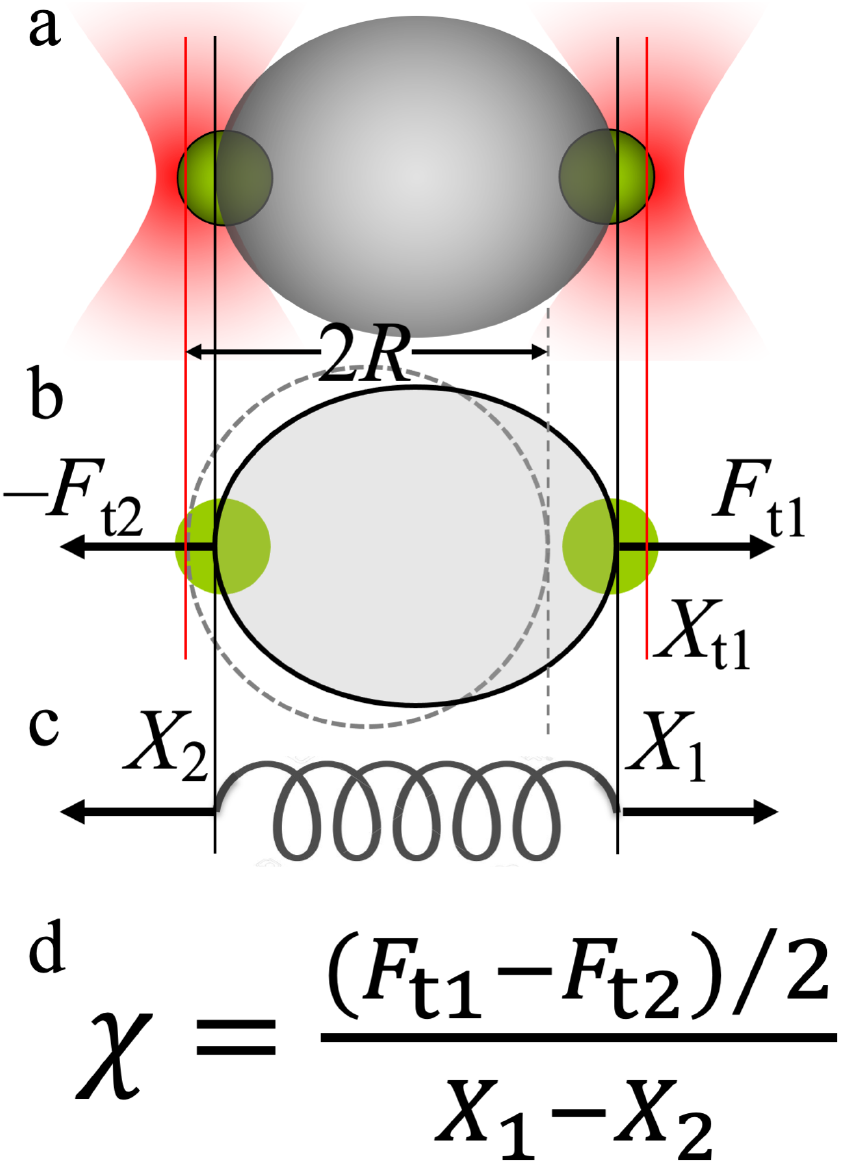
Experimental setup for determining the effective spring constant *χ*(*ω*) of biomolecular droplets. (a) Side view: two optically trapped polystyrene beads pull at the poles of a droplet. Trap 2 is fixed at the origin while trap 1’s position undergoes harmonic oscillation. (b) Top view: ellipse (circle) with solid (dashed) boundary shows the deformed (unperturbed) droplet. *F*_t1_ and *F*_t2_ are the forces exerted by the traps on the beads; *X*_t1_ is the position of trap 1; *X*_1_ and *X*_2_ are the positions of the beads. (c) The deformation of the droplet is equivalent to the oscillation of a spring. (d) The effective spring constant.

However, with actual experimental *χ*(*ω*) data, the extraction of *G*\*(*ω*) faces serious problems. The first problem is that, without an independently determined value for the surface tension, *χ*(*ω*) data alone are insufficient to extract both *γ* and *G*\*(*ω*). Both *χ*(*ω*) and *G*\*(*ω*) have real and imaginary parts. In essence one is trying to determine three quantities [*γ*, *G*′(*ω*), and *G*″(*ω*)] from two knowns [*χ*′(*ω*) and *χ*"(*ω*), denoting the real and imaginary parts of *χ*(*ω*)]. Jawerth et al. used the average of *χ*′(*ω*) at the lowest three frequencies studied (*ω*/2*π* at 0.04, 0.1, and 0.2 Hz) to set *γ*, assuming negligible contribution from *G*\*(*ω*) at such low frequencies. Following this recipe and using Eq. (16), I obtain *γ* = 6.2 μN/m for a set of *χ*(*ω*) data reported by Jawerth et al. at 75 mM KCl [presented in their Fig. 3(a)]. However, *γ* cannot be uniquely determined. For a given *γ*, a *G*\*(*ω*) can be found that together with *γ* reproduces the *χ*(*ω*) data according to Eq. (16). Figure 2(a) presents *G*\*(*ω*) results corresponding to four different *γ* values: 6.2, 4.91, 4.0, and 0 μN/m; all the four *γ*−*G*\*(*ω*) combinations, including the extreme case with 0 surface tension, reproduce the same set of *χ*(*ω*) data, which is reproduced in Fig. 2(b) as circles.

**FIG. 2.**
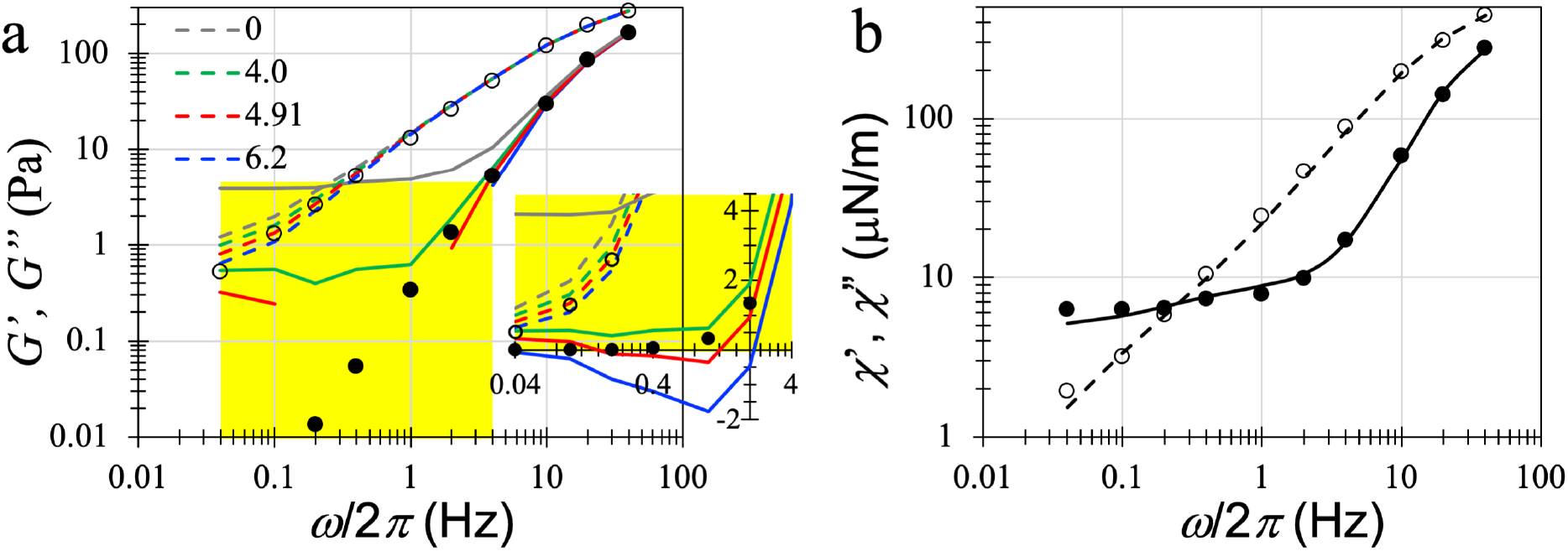
Problems with extraction of *γ* and *G*\*(*ω*) and a solution. (a) Four sets of *G*\*(*ω*), corresponding to four choices of *γ*, all reproduce the same set of *χ*(*ω*) data. Extraction of *G*\*(*ω*) at a given *γ* is according to Ref. 14, with *l* = 40,000. Solid and dashed curves display real and imaginary parts of *G*\*(*ω*), respectively; the legend displays the *γ* values. Pieces of *G*′(*ω*) missing on the ordinance log scale (yellow shading) correspond to negative values, which are shown on a linear scale in the inset. Filled and open circles display the Jeffreys model for *G*′(*ω*) and *G*″(*ω*). (b) The *χ*(*ω*) data of Jawerth et al. at 75 mM KCl, shown as filled and open circles, are accurately reproduced by the Jeffreys model (curves).

**FIG. 3.**
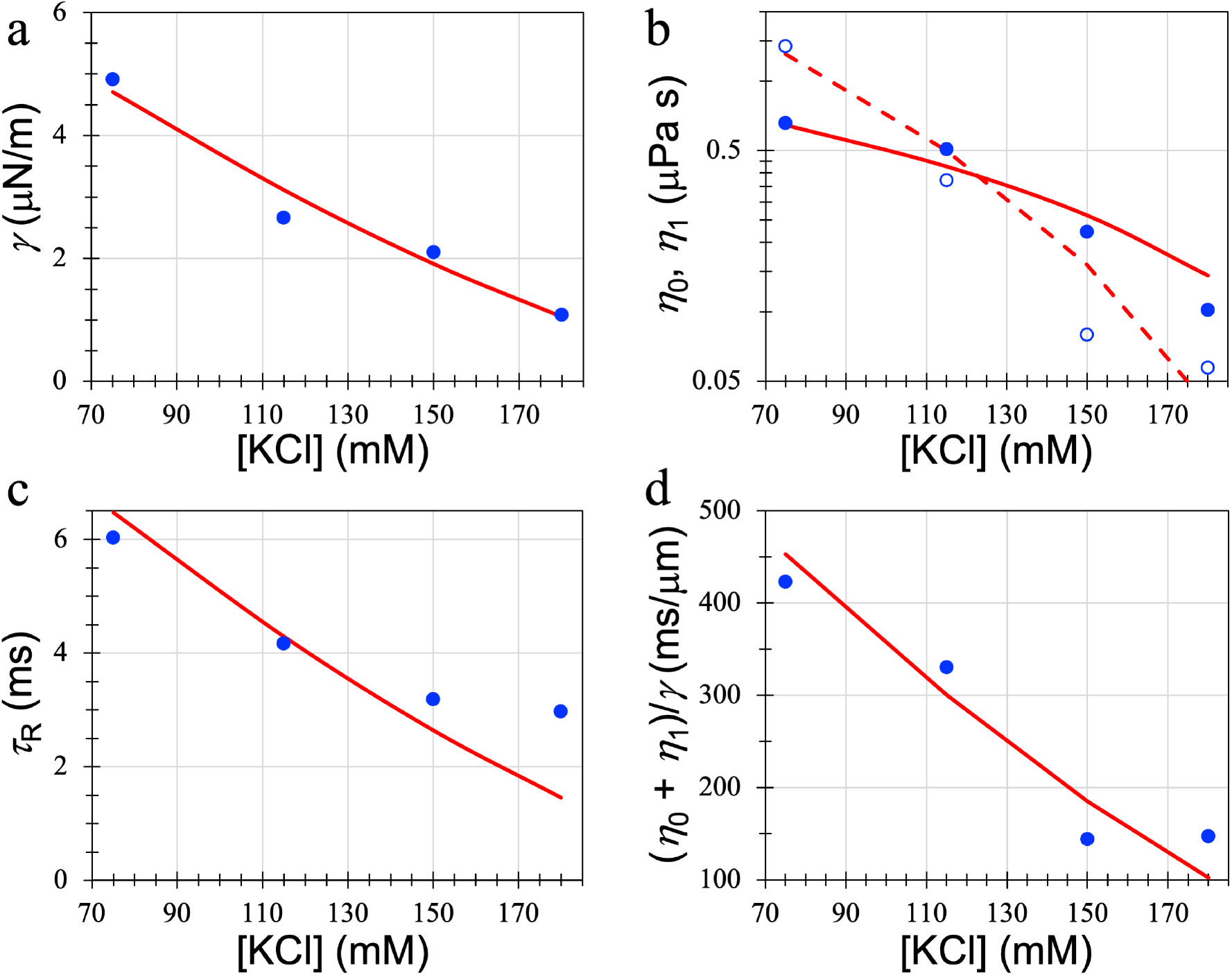
Salt dependences of surface tension and the three parameters in the Jeffreys model of viscoelasticity. Circles display data obtained from the fitting procedure to the *χ*(*ω*) data of Jawerth et al. at four KCl concentrations; curves are fits to a scaling law, *γ* = *A*([KCl]_c_ − [KCl])α; [KCl]_c_ = 241 mM and *α* is set to 3/2, except for η_1_. (a) *γ*. (b) η_0_ (filled circles and solid curve) η_1_ (open circles and dashed curve). For η_1_ the critical exponent is increased to 3.5. (c) *τ*_R_. (d) (η_0_ + η_1_)/*γ*.

The second problem in the extraction of *G*\*(*ω*) is that some of the extracted *G*′(*ω*) values at low frequencies are negative [see Fig. 2(a) inset], and hence unphysical. For example, with *γ* = 6.2 μN/m, *G*′(*ω*) is −1.8 Pa at 1 Hz. The negative *G*′(*ω*) values might be taken as an indication that the choice of *γ* is too high, as *G*′(*ω*) is no longer negative when *γ* is reduced to 4.0 μN/m. However, the deeper reason is that *G*′(*ω*) at low frequencies is particularly sensitive to errors in the *χ*(*ω*) data, as illustrated by the large differences of the low-frequency *G*′(*ω*) values extracted with the three non-zero *γ* choices. As *G*′(*ω*) is expected to approach 0 rapidly at low frequencies, errors in *χ*(*ω*) may inevitably lead to negative values for the extracted *G*′(*ω*).

To solve both of these problems, I propose choosing an appropriate model for *G*\*(*ω*) and using the corresponding *χ*(*ω*) to fit experimental data. Ref. 14 already gave an indication that the shape of *χ*(*ω*) produced by the Jeffreys model looks similar to the experimental data of Jawerth et al. [circles in Fig. 2(b)]. By scanning over the values of four parameters: *γ*, η_0_, η_1_, and *τ*_R_, the *χ*(*ω*) data of Jawerth et al. can be accurately reproduced [curves in Fig. 2(b)], with *γ* = 4.91 μN/m, η_0_ = 0.659 Pa s, η_1_ = 1.42 Pa s, and *τ*_R_ = 6.03 ms. This solution is unique as all other solutions lead to larger deviations from the experimental data for *χ*(*ω*). Positivity of *G*′(*ω*) is guaranteed in the Jeffreys model. The *G*\*(*ω*) with the above parameter values is displayed as circles in Fig. 2(a). It is closest to the *G*\*(*ω*) [red curves in Fig. 2(a)] extracted from the *χ*(*ω*) data by the numerical procedure when *γ* is set to the optimal fitting value, i.e., 4.91 μN/m, but avoids the negative values of the extracted *G*′(*ω*). Our preliminary experimental data for the viscoelasticity of a variety of biomolecular droplets also appear to fit the Jeffreys model (Ghosh and Zhou, unpublished).

Applying the same fitting procedure to the *χ*(*ω*) data of Jawerth et al. for PGL-3 droplets at other KCl concentrations, I obtain the salt dependences of the surface tension and the parameters, η_0_, η_1_, and *τ*_R_, of the Jeffreys model. Figure 3(a) displays the *γ* values at four KCl concentrations: 75, 115, 150, and 180 mM. *γ* spans the range of 1 to 5 μN/m and decreases with increasing salt concentration, as expected from the salt screening of electrostatic attraction between PGL-3 molecules inside droplets. The salt dependence can be represented by a scaling law *γ* = *A*([KCl]_c_ − [KCl])^α,16^ with *α* = 3/2 as the critical exponent and [KCl]_c_ = 241 mM as the critical salt concentration. η_0_ decreases from 0.66 to 0.10 Pa s, and the salt dependence can be fit to the same scaling law [solid circles and solid curve in Fig. 3(b)]. Meanwhile, η_1_ decreases from 1.42 to 0.057 Pa s and thus shows a much stronger salt dependence, requiring a rise of the critical exponent to 3.5 [open circles and dashed curve in Fig. 3(b)]. As a result of their disparate salt dependences, the viscosity of PGL-3 droplets switches from being dominated by η_1_ at low salt to being dominated by η_0_ at high salt. In other words, the droplets move closer to being a Newtonian fluid at high salt. While viscosity is expected to decrease with increasing salt concentration due to weakened electrostatic attraction, there is no simple explanation why the Newtonian component and Maxwellian component show different salt sensitivities. The shear relaxation time constant *τ*_R_ decreases from 6.03 to 3.0 ms, and the salt dependence can also be represented by the scaling law with the regular critical exponent 3/2 [Fig. 3(c)].

The dynamics of the shape change of biomolecular droplets depends on their surface tension and viscosity, with the former driving the relaxation toward a spherical shape while latter impedes the process. The combination, η*R*/*γ*, determines the time scale of the dynamics. In particular, the fusion time for two droplets with an equal radius *R* is τ_f_ = 1.97η*R*/*γ*.^12^ For the Jeffreys model, we expect (η_0_ + η_1_)*R*/*γ* to play a similar role. For PGL-3 droplets, the inverse speed, (η_0_ + η_1_)/*γ*, decreases from 423 ms/μm at 75 mM KCl to 147 ms/μm at 180 mM KCl, and the salt dependence also fits the scaling law with the regular critical exponent 3/2 [Fig. 3(d)]. At *R* = 5 μm, the predicted fusion times decrease from 4170 ms at 75 mM KCl to 1450 ms at 180 mM KCl. Three comments are in order. First, the predicted fusion times can be directly tested by experiments. Second, these fusion times are three orders of magnitude longer than the corresponding shear relaxation times. Therefore, during the fusion of PGL-3 droplets, shear relaxation inside the droplets is effectively instantaneous and the droplets can be treated as a Newtonian fluid, justifying such treatment in previous studies^12^ and the present use of (η_0_ + η_1_)*R*/*γ* as the time scale of droplet shape dynamics when invoking the Jeffreys model. Third, the predicted 3-fold speedup in PGL-3 droplet fusion dynamics is due to the different rates at which surface tension and viscosity decrease with increasing KCl concentration. The decrease in viscosity occurs at a higher rate and, as noted above [Fig. 3(b)], this is especially true for the Maxwellian component. A 4-fold speedup in fusion dynamics was actually observed for droplets formed by the binary mixture a pentameric construct of proline-rich motifs and a polymer, heparin, upon increasing KCl concentration from 150 mM to 400 mM.^12^

The fusion times of different biomolecular droplets span several orders of magnitudes.^12^ For example, droplets formed by the mixture of polylysine and heparin fuse within a few ms, whereas the fusion of droplets formed by the mixture of lysozyme and a pentameric construct of SH3 domains takes as long as 10,000 ms. The slow-fusion droplets, similar to PGL-3 droplets, may behave viscoelastically as Newtonian fluids. However, for the fast-fusion droplets, shear relaxation may become rate-limiting for the fusion process. Our preliminary experimental data suggest that polylysine:heparin droplets may present precisely such a case (Ghosh and Zhou, unpublished).

The biology of biomolecular condensates as a result of recent intense studies is far outstripping the physical understanding of the underlying material properties. The conformality to the Jeffreys model of viscoelasticity demonstrated here will facilitate both the experimental determination of viscoelastic properties and the comparison of these properties at different solvent conditions and among different condensates. The present work helps with mechanistic interpretation regarding the dynamics of biomolecular condensates, but also raise even deeper questions on the parameters of the Jeffreys model. In particular, what are the molecular motions that dominate shear relaxation, which can potentially rate-limit droplet fusion? Progress in these directions will narrow the gap in understanding between the biology and physics of biomolecular condensates.

## DATA AVAILABILITY

The data that support the findings of this study are available from the corresponding author upon reasonable request.

## ACKNOWLEDGMENTS

This work was supported by National Institutes of Health Grant GM118091.

